# Evidence for Transcriptomic Conservation Between the Main Cells of the *Drosophila* Prostate-Like Accessory Gland and Basal Cells of the Mammalian Prostate

**DOI:** 10.1101/2025.06.05.658085

**Authors:** S. Jaimian Church, April R. Kriebel, Joshua D. Welch, Laura A. Buttitta

## Abstract

The *Drosophila* accessory gland performs functions analogous to the mammalian prostate in production of seminal fluid components that are essential for male fertility. The mammalian prostate and *Drosophila* accessory glands share a similar tissue organization and structure. Both organs contain secretory epithelial cells forming a gland lumen, surrounded by a stroma with extracellular matrix enveloped by innervated muscle for organ contraction and fluid release. However, the *Drosophila* accessory gland secretory epithelium is postmitotic, polyploid and binucleate, and lacks a known stem cell population. By contrast, the mammalian prostate epithelium is made up of diploid luminal secretory cells and diploid basal cells that are maintained by at least two stem cell populations. Despite the differences in the tissues, it has been argued these tissues may share a ‘deep homology’ based on the expression of conserved genes during development. Here we performed a cross-species comparative analysis using single-cell RNA sequencing data from adult tissues using data from the *Drosophila* Fly Cell Atlas and mammalian adult prostate single-cell datasets. Our analysis provides additional evidence of transcriptomic similarity between the main epithelial cells of the *Drosophila* prostate-like accessory gland and the basal epithelial cells of the mammalian prostate. While we do not know whether these similarities reflect shared evolutionary homology, or independently derived features due to shared tissue functions, our results strengthen the arguments that the *Drosophila* accessory gland can be used to effectively model aspects of human prostate biology and disease.

## Introduction

We have learned about fundamental biological processes by studying a few model organisms that are well-suited for experimental research. These organisms reveal insights into cellular pathways, developmental processes and organ morphogenesis and homeostasis conserved across millions of years of evolution. *Drosophila melanogaster*, the fruit fly, stands out among these models for contributing to our understanding of pathways involved in developmental tissue patterning and differentiation. For over a century, the fruit fly has been a cornerstone in studies of genetics, development, aging, and physiology (Ashburner 1989; Chapman et al. 1995; Morgan 1910; Nusslein-Volhard and Wieschaus 1980). Despite the ∼800 million years of evolution between invertebrates and vertebrates (Hedges et al. 2006), several genetic networks and pathways perform similar functions in flies and mammals, with examples including the eye developmental regulatory network (Callaerts et al. 1997), roles for the Toll signaling pathway in innate immunity (Lemaitre et al. 1996), and tissue growth and size control via the Hippo pathway (Dong et al. 2007; Staley and Irvine 2012). Deeper examinations of gene regulatory networks used during development have revealed unexpected conservation of networks underlying ‘deep homology’ and a potentially shared origin of anatomical structures that appear superficially variable and quite different (Shubin et al. 2009, 1997). The level of deep homology in developmental gene regulatory networks is further emphasized in examinations of discrete genetic regulatory elements (Wong et al. 2020), and the evolution of tissue-specific gene expression (Mantica et al. 2024). With the advance of single-cell transcriptomics and multi-omics, researchers have unprecedented resolution to perform cross-species modeling on biological datasets from complex tissues and organs at various stages of development and adulthood. Performing cross-species comparisons on datasets can offer deeper insights into shared and unique genes or pathways, enhancing our understanding of biology throughout evolution (Bakken et al. 2021; Musser et al. 2021; Woych et al. 2022; Yoon et al. 2023).

The *Drosophila* male accessory gland is a tissue functionally analogous to the mammalian prostate (Ito et al. 2014; Rambur et al. 2021; Wilson et al. 2017). These organs share some similarities in the seminal fluid components they produce for reproductive success and their cellular mechanisms for secretion and exosome release (Garenaux et al. 2015; Kratzschmar et al. 1996; Ram and Wolfner 2009; Udby et al. 2005; Wilson et al. 2017), but also have distinct, species specific products. One product is the *Drosophila* protein Sex Peptide (SP), which mediates physiological changes in *Drosophila* females after mating (Chapman et al. 2003; Chen et al. 1988; Liu and Kubli 2003; Tsuda and Aigaki 2016). The organ architecture is broadly similar, with both the mammalian and *Drosophila* tissues containing secretory epithelial cells forming a lumen surrounded by a stroma or extracellular matrix, enveloped by an innervated muscle layer to aid in tissue contraction and fluid release (Bairati 1968; Shen and Abate-Shen 2010; Wang et al. 2018; Wilson et al. 2017). However, there are also significant differences in the tissue organization and development between insects and mammals. The mammalian prostate epithelium contains three cell types, basal cells located next to the basal lamina, luminal cells located more apically, and rarer neuroendocrine cells interspersed in the luminal cell layer. In the *Drosophila* tissue, there are two known secretory epithelial cell types organized in a monolayer, with one cell type, the main cells making up the majority (∼1000 cells) of each gland and a larger, highly secretory cell type, termed secondary cells comprising 40-60 cells per gland.

Detailed single-cell transcriptomics of the accessory glands across multiple species have suggested there may be additional subtypes of cells within the *Drosophila* main cell population, but molecular markers to distinguish these main cell subtypes are still lacking (Majane et al. 2022). Unlike the diploid epithelial cell types of the mammalian organ, the main and secondary cells of the *Drosophila* tissue are unique in that they are binucleate and polyploid. The binucleate and polyploid properties of these cell types is due to a terminal differentiation program during development that involves cell cycle variants that successively truncate the final cell cycles. This results in cells with two octoploid nuclei in the mature adult gland (Box et al. 2024; Taniguchi et al. 2018; Taniguchi et al. 2014; Kiichiro Taniguchi 2012). In addition, there is no known stem cell population for the adult *Drosophila* organ, which is thought to be entirely postmitotic. In contrast, stem cell populations for the mammalian organs have been identified, allowing for robust tissue regeneration (Li and Shen 2019). Finally, the mammalian tissues develop from the endodermal layer of the embryo, while the *Drosophila* accessory glands develop from mesoderm (Ahmad and Baker 2002), further highlighting important differences in the development of these organs.

Despite their shared functions, the anatomical and developmental differences between the mammalian prostate and *Drosophila* accessory gland have led to these tissues usually being described as analogous rather than homologous. However, studies of accessory gland development have revealed evidence for ‘deep homology’ between the insect and mammalian organs, with evidence of a common developmental program using conserved FGF signaling pathways involved in tubulogenesis and Hedgehog and BMP pathways in patterning organ morphogenesis (Kumari and Sinha 2021). While the developmental programs of prostate-like organ formation may reveal some deep homology, we wondered whether detailed, cell-type specific transcriptomic analysis of the terminally differentiated adult tissues could reveal shared gene expression features and potentially illuminate additional shared gene regulatory networks of specific cell types. For example, are *Drosophila* main cells potentially more similar to mammalian basal or luminal cells?

To compare cell-type specific gene expression in adult tissues, we took advantage of the recently published Fly Cell Atlas (Li et al. 2022), a single-cell transcriptomic atlas containing many adult *Drosophila* organs, including those of the male reproductive system. We compared the Fly Cell Atlas male reproductive system data to a high-quality mouse single-cell prostate dataset, GSM4556594 (Crowley et al. 2020). Cross-species comparisons between mouse and human prostate cells have been made, but comparisons across further evolutionary distances are challenging (Hedges 2002). Two rounds of whole genome duplication have occurred between these species (Dehal and Boore 2005), leading to a more complicated genetic network in mammals. In addition, integrating single-cell (sc)RNA-seq datasets is challenging due to batch effects, variations in cell type composition, sequencing depth differences, and the heterogeneity of cell populations (Camara 2018; Luecken et al. 2022). Overcoming these obstacles requires advanced computational methods capable of normalizing data, countering batch effects, and precisely matching cell types and states while retaining each dataset’s distinct biological insights. In the realm of single-cell data integration, various tools have been developed to address the challenges of comparing and integrating datasets from different experimental setups. LIGER, Harmony, and Seurat are prominent examples of such tools, each with its own approach and strengths (Chazarra-Gil et al. 2021; Luecken et al. 2022; Tran et al. 2020). LIGER stands out as an option for cross-species comparison due to its nonnegative factorization methodology, which processes gene expression data to uncover shared and unique features from multiple datasets (Liu et al. 2020; Welch et al. 2019). This approach facilitates the identification of common gene expression features across different datasets while highlighting dataset-specific factors, effectively retaining the unique aspects of each biological sample (Liu et al. 2020). Other methods, like Harmony take a different approach with its multi-dataset alignment algorithm, which is adept at identifying shared cell states and correcting batch effects (Korsunsky et al. 2019), which may work best for datasets from similar conditions or closely related species (Korsunsky et al. 2019; DePasquale et al. 2019). Seurat also employs an integration strategy that seeks "anchors" between datasets to mitigate batch effects and align similar cell types (Stuart et al. 2019). While Seurat is widely recognized for its effectiveness in integrating and analyzing single-cell data, its approach emphasizes the identification of shared cellular states, possibly at the expense of species-specific or condition-specific features that differentiate one dataset from another.

In this study we used LIGER to compare the *Drosophila* male reproductive system to the mouse prostate. We found significant gene expression similarities that map the *Drosophila* accessory gland main cells to the basal cells of the mouse prostate. Importantly, when we compare other *Drosophila* tissues, including secretory epithelium such as the intestine, we do not see such overlap, suggesting the gene expression similarities we uncover between the *Drosophila* main cells and mouse basal cells may be due to true shared tissue and cell type-specific expression patterns.

## Results and Discussion

The first challenge for our cross-species comparison was to find the appropriate *Drosophila* to mammalian orthologs (Figure 1A). Our study utilized the ortholog prediction tool DIOPT (www.flyrnai.org/cgi-bin/DRSC_orthologs.pl) to determine orthologs between *Drosophila* and humans or mice and humans. Due to genome-wide duplications between invertebrates and vertebrates, as well as gene specific duplications leading to multiple paralogs, DIOPT’s predictions include numerous duplicates for many genes, as shown in Figure 1. The largest category of gene pairs is one-to-one gene pairing, with about 8,000 genes in this category.

**Figure 1:**
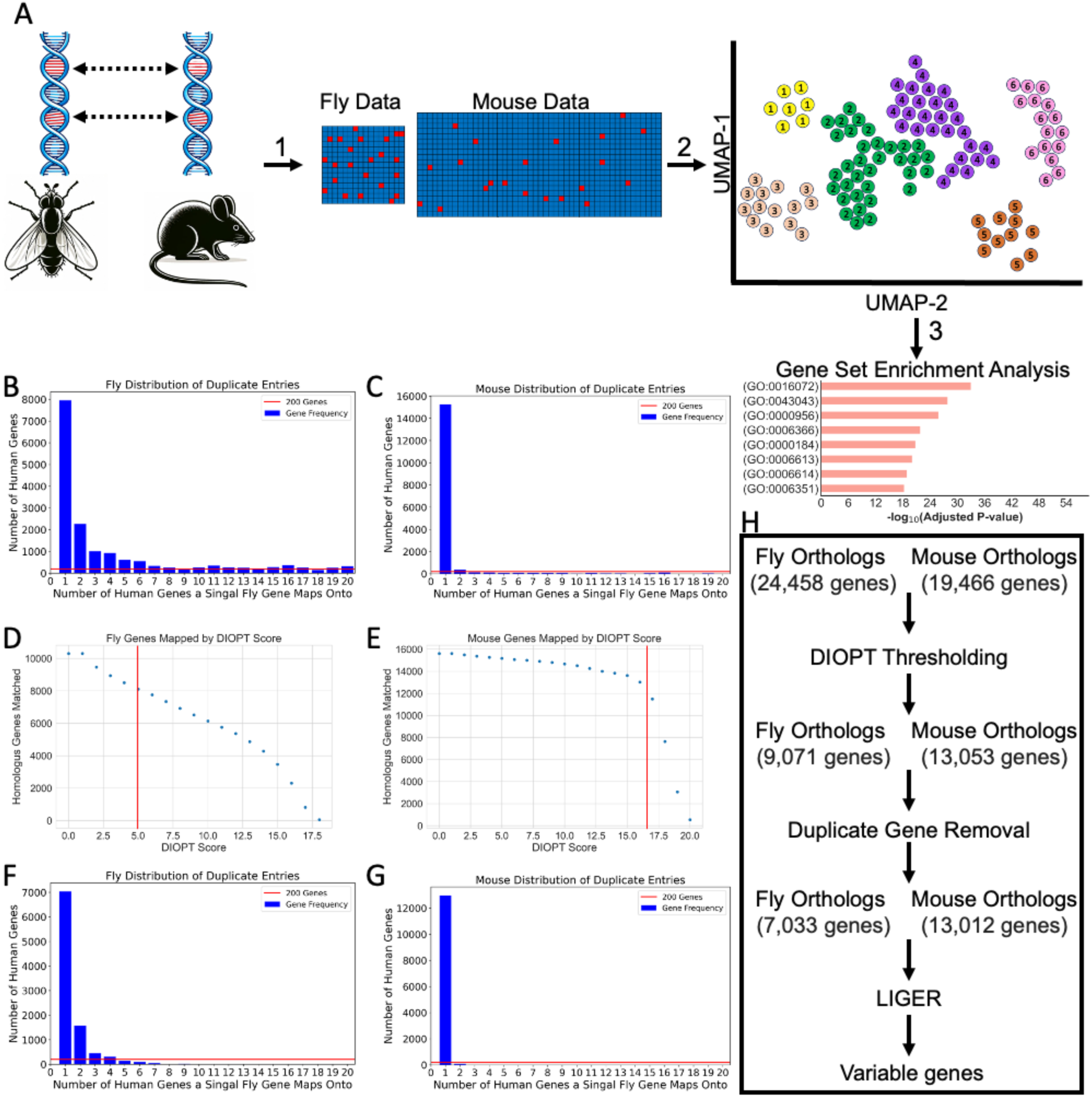
DIOPT Filtering of Low Scoring predicted Orthologous Genes between *Drosophil*a and Human Result in 7,000 One-to-One Gene Matching Pairs A) Schematic representation of how LIGER can be used to analyze biological datasets from different species to find shared features. B,C) Numbers of duplicated genes in the fly and mouse prostate datasets. D,E) Scatterplots of the DIOPT scores for the fly and mouse datasets compared to human. F,G) Numbers of genes with more than one match to human genes in the fly and mouse prostate datasets after thresholding the datasets on DIOPT scores indicated by the red lines in D and E.

However, we can see on the tail end of the graph that there are a small number of fly genes with 20 or more possible human orthologs (Figure 1 B). While we capped Figure 1, panels B and C at 20 possible human orthologs, both datasets had gene pairs that even surpassed 20 possible orthologs. The distribution in Figure 1B suggests that many of the duplicated genes arise from a small number of fly genes with a large number of possible human orthologs, rather than most fly genes having 2 or 3 possible human orthologs. However, fly genes with 5 or more orthologs comprised more than half of all the fly-to-human orthologous pairs (12,303 human genes), illustrating the complexity of such an evolutionarily distant cross-species comparison. By contrast, 15,263 genes of the 17,822 mouse-to-human possible orthologs were one-to-one matches.

We next examined whether there may be a pattern in the accumulated DIOPT score for predicted matches that could indicate the quality of homolog mating. Identifying a pattern would allow us to create a cutoff score for selecting the most probable one-to-match for homolog mating, consequently increasing the number of genes that could be treated as one-to-one matches. We plotted DIOPT scores for both datasets, comparing fly to human genes and mouse to human genes (Figures 1 D and E). For the mouse to human comparison, we find a DIOPT score threshold where the relationship between DIOPT scores and the number of matches diverged. At a DIOPT score of 16, there was a clear bifurcation with scores below 16 allowing many weak matches with duplicates, but without significantly increasing the total number of unique human genes with matches. This gave us a data-based threshold for the DIOPT score, allowing us to optimize the number of genes represented while retaining only strong matches. By imposing a DIOPT threshold, many weak gene pairs and duplicate matches with weaker scores were removed. In contrast, the fly dataset showed a more linear decrease in the number of genes with matches (y-axis) as the DIOPT score increased (x-axis, Fig. 1D). Unlike the mouse-human comparison, there was no clear DIOPT threshold where the relationship between DIOPT scores and the number of genes diverged. This made it much more challenging to impose a DIOPT score threshold to remove weak and duplicate matches. After manually examining dozens of genes for which orthologous functions are known, we elected to impose a DIOPT score cutoff of 5, hoping to retain true orthologous matches, but minimize weak or duplicate matches. This issue highlights one of the challenges of distant cross-species comparisons.

After imposing our DIOPT score thresholds, 9,071 gene pairs for fly-to-human comparisons remained, with 7,033 of these fly genes exhibiting one-to-one matches which were used for downstream analysis. For the mouse-to-human orthologs, 13,053 predicted one-to-one matching orthologs were used for downstream analysis. While we recognize that many developmentally and evolutionarily important genes are removed using these thresholds and cutoffs, we decided to proceed with cross-species comparisons using these high confidence one-to-one orthologous gene sets (Fig. 1H).

The next challenge was to annotate the cells within our mouse and fly datasets. The cells in the Fly Cell Atlas male reproductive system dataset had annotations provided, which we utilized (Li et al., 2022). The mouse prostate dataset did not have annotation data integrated, or a list of marker genes used for their methodology (Crowley et al., 2020). In mice, the prostate organ has four lobes, anterior, dorsal, ventral, and lateral, encircling the urethra rather than a singular anatomical structure (Oliveira et al. 2016). We used a literature search of common gene markers and the Human Protein Atlas (https://www.proteinatlas.org/) to annotate mouse prostate cell types (Crowley et al. 2020; Aparicio et al. 2025; Li et al. 2024). Because our focus was determining the similarities and differences between mouse basal and luminal cells to compare to the fly epithelial cell types, we did not seek to separate the mouse prostate cells into as many divergent populations as possible. We used the Python ScanPy pipeline (Wolf et al. 2018) to normalize and cluster the mouse prostate cells (Figure 2A). We annotated the cluster from Figure 2A and Figure 2B using a set of 54 genes manually curated from the mouse prostate literature.

**Figure 2:**
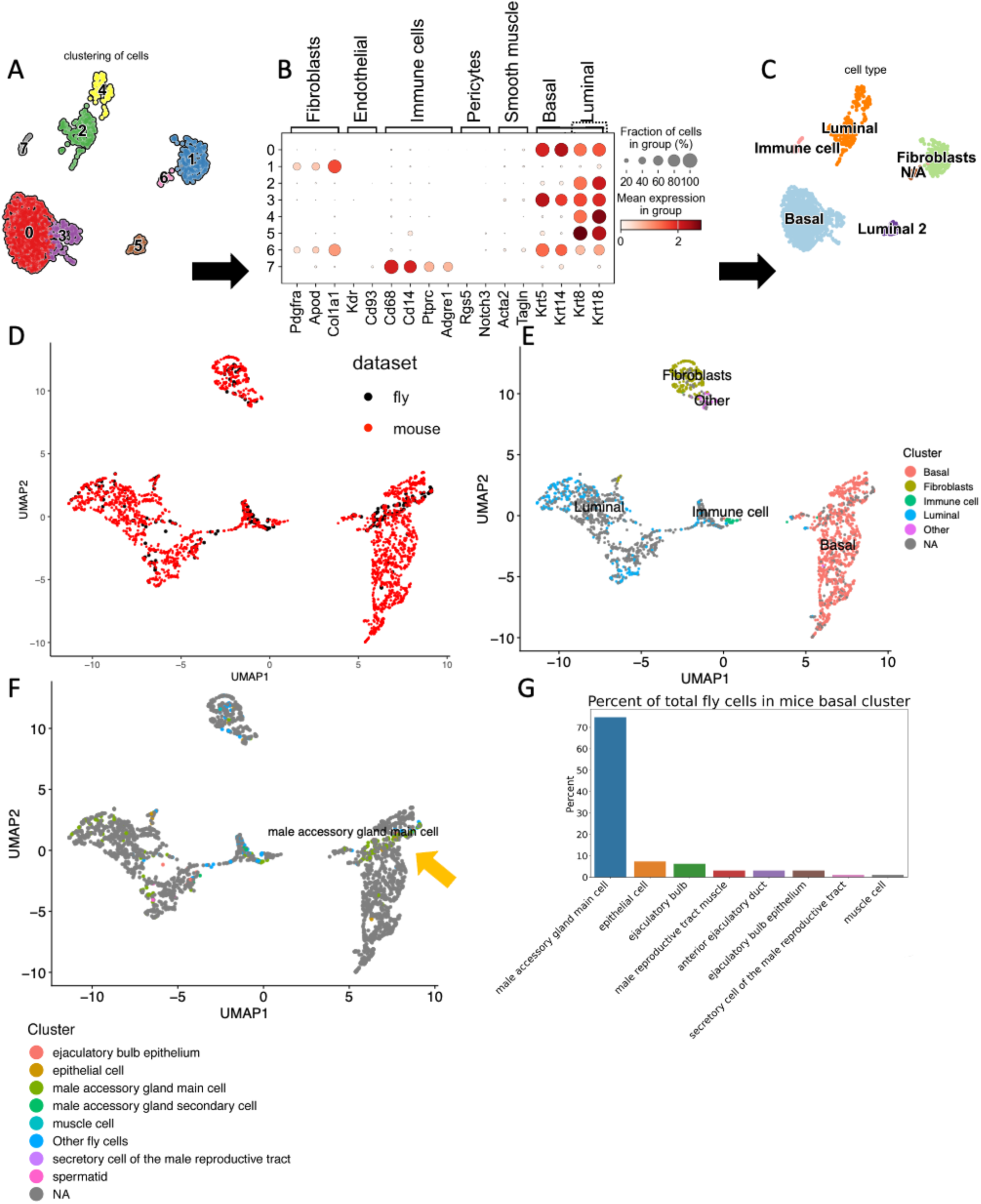
D*r*osophila Male Reproductive System Cells Cluster with Mouse Prostate Cells A) Mouse cells were processed via ScanPY’s pipeline using Leiden clustering. B) Marker genes used for annotation of mouse prostate cell clusters. C) Annotated cell types from panel A. D) UMAP plot of fly male reproductive system mapped onto mouse prostate cells after LIGER integration performed using only shared genes. E-F) UMAP plots from panel D with the mouse or fly cell annotations overlaid. The yellow arrow indicates fly main cells in the mouse basal population. G) The distribution of fly cells mapping to the mouse prostate basal cell cluster.

We found 17 of these mouse prostate cell markers to be informative for cell type clustering. See Supplementary Figure 1 for the complete list of genes tested for annotating the clusters. The finalized cluster annotations are displayed in Figure 2C, revealing one basal mouse population, two luminal populations, a fibroblast population, an immune cell population, and a small group of unclassified cells labeled as “NA” that did not align with any known cell types in the mouse prostate based upon our molecular markers.

In the process of integrating multiple datasets using LIGER, setting a reference dataset is a crucial step that serves as a foundational basis for the integration. The reference dataset acts as an anchor, providing a standard or baseline against which other datasets can be aligned and compared. This is particularly important in studies where cross-species comparisons are made, as the datasets are often different sizes, and the biological variability can be significant. By establishing a reference, LIGER can more effectively identify shared cellular states and features across the datasets, ensuring the integration is grounded in a consistent biological context. The choice of the reference dataset in LIGER should be the most robust and biologically relevant dataset. The mouse dataset had the greatest number of cells and greater gene coverage within cells. As such, we selected the mouse prostate dataset as our reference. Figure 2D shows the integration of the fly and mouse datasets. The fly cells cluster together with the mouse cells, indicating that these cells share patterns of gene expression (Figure 2D). In Figure 2E, we can see that LIGER has separated the cels into the cell type clusters that were previously identified in Figure 2C with Scanpy. Next, we looked at the purity scores for the mouse reference dataset. The purity score is an indicator of how well the clustering reflects the original clustering and annotation of a dataset. To calculate cluster purity, we annotate each cluster in the shared latent space and the cells within that cluster. Using the original annotation labels, we then calculate the percentage of cells that still retain their original annotations despite being re-annotated within the new latent space. For the UMAPs seen in Figures 2D-2F, the mouse purity score was 97%, indicating that 97% of cells annotated as a particular cell type were again identified as that cell type in our LIGER integration. In particular, the mouse basal population, which we focus on later, had a purity score above 99%.

With fly cells mapping onto mouse clusters, we next need to determine what our comparison would call significant clustering vs nonsignificant clustering. As our focus is identifying cell-type specific shared gene expression, fly cell types should be enriched and have high purity to have significant co-clustering onto mouse clusters. To determine enriched thresholds of a cluster, a fly cell type must have more than 50% of cells in the cluster in question. To determine the purity threshold within a cluster, the fly cell type must comprise more than 50% of the cells within the cluster. Using these thresholds, we compared the cells of the fly reproductive system to that of the mouse prostate, seen to have co-clustering Figure 2D, F. Only the male accessory gland main cells met our threshold for enrichment, with 55% of all main cells clustering within the mouse basal populations. The other annotated cell types in the reproductive system dataset were sparsely represented, with several cell types having fewer than 10 annotated cells. We, therefore, chose to focus solely on the male accessory gland main cells for further analysis.

The purity of the male accessory gland main cells mapping onto the mouse basal cell cluster was also high, as we found the main cells to comprise nearly ∼75% of all the fly cells within this cluster Figure 2G. None of the other fly cell types mapping onto the mouse basal population reached higher than ten percent (Figure 2G). This led us to wonder whether the high purity and positive mapping onto this cluster may be an artifact due to the relatively large population size for the main cells in the *Drosophila* dataset. To address this, we also compared *Drosophila* main cells to cells from other mammalian tissues using the Tabula Muris Senis Mouse Aging Cell Atlas, a comprehensive compilation of single-cell transcriptomic data spanning various tissues and organs from mice at different life stages (Tabula Muris et al. 2018). We performed a comparison between the *Drosophila* male accessory gland main cells and mouse intestinal epithelial secretory cells, again using the mouse dataset as the reference dataset. We observed little co-clustering of the *Drosophila* main cells with the mouse intestine (Fig 3B) suggesting there is little spurious overlap with datasets containing secretory epithelial cells, a finding we further confirmed by testing for overlap with the highly secretory mouse acinar cells of the pancreas. By contrast, we observed significant overlap and enrichment for clustering when comparing fly intestine to mouse intestine, providing confidence that we can identify true cell-type specific similarities (Fig. 3A).

**Figure 3:**
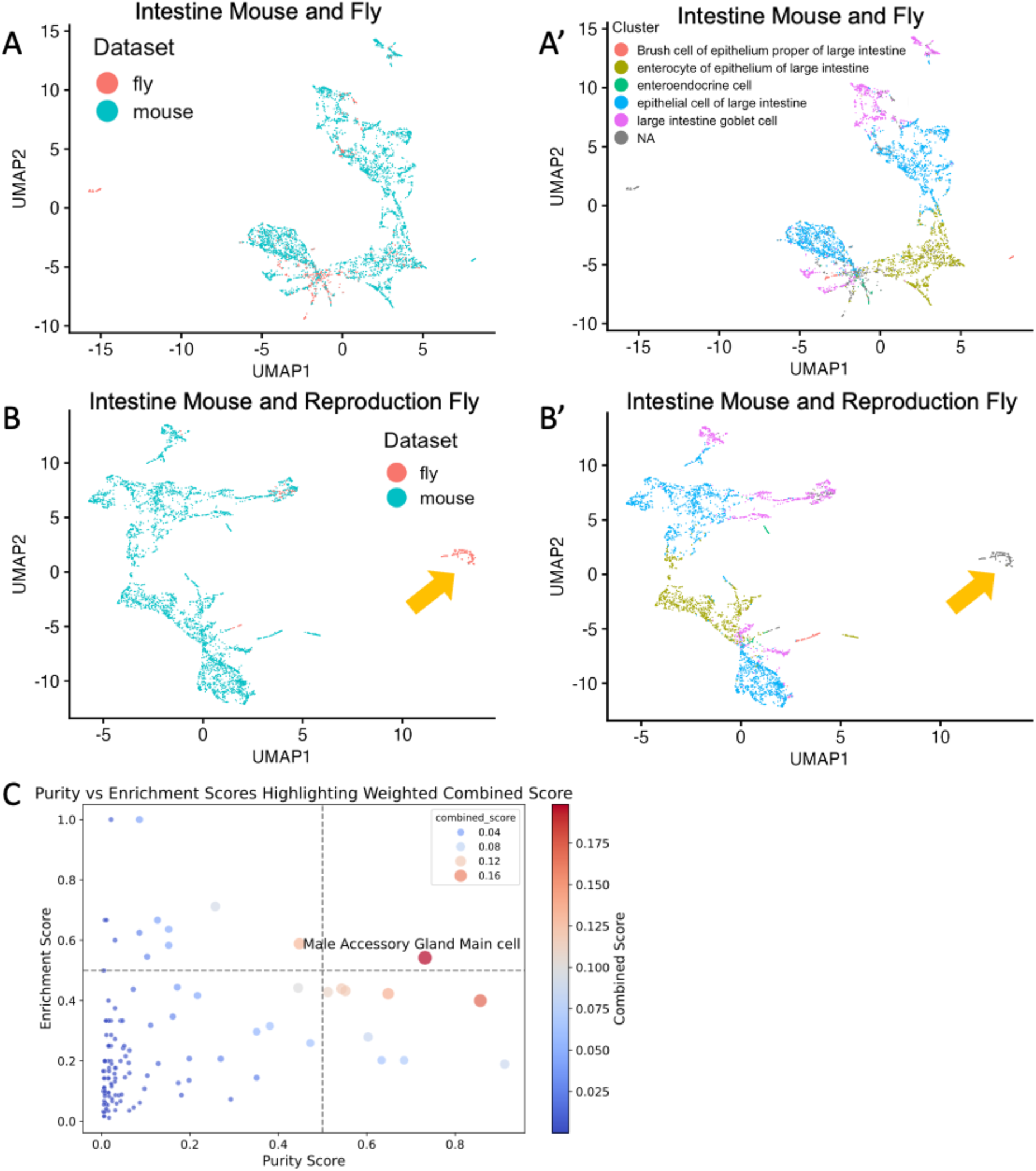
D*r*osophila Accessory Gland Main Cells Show Significant Clustering with the Mouse Prostate Basal Cell Population A) UMAP plots of fly and mouse intestinal cells clustered after LIGER integration performed using only shared genes. A’) Shows the mouse annotation for panel A. B) UMAP plots for fly male reproductive cells and mouse intestinal cells clustering after LIGER integration performed using only shared genes. B’) Shows the mouse annotation for panel B. The yellow arrow indicates fly reproductive cells clustering away from the mouse cell populations. C) Plot showing purity vs enrichment scores for 212 fly cell types mapped onto the mouse basal cell cluster. The X-axis shows the purity of an individual fly cell in the mouse basal populations. The Y-axis shows the enrichment of individual fly cells in the mouse basal populations. Taking the product of these values, we get a combined score. The combined score is indicated by dot color and size. Of all 212 fly cell types, the accessory gland main cells have the best combined score match to the mouse prostate basal cells.

We next tested whether any other cell type in *Drosophila* can map onto the mouse basal cell cluster, as we see for main cells. To do this, we plotted the purity vs. enrichment scores for 212 *Drosophila* cell types from nearly all adult tissues present in the Fly Cell Atlas, when mapped onto the mouse prostate basal populations as in Fig. 2F. A fly cell type was considered to co-cluster with the mouse basal cluster if its purity and enrichment scores were 0.5 or above. Of the 212 *Drosophila* cell types tested, 125 cell types had at least one cell map onto the mouse basal population Figure 3C, but only the male accessory gland main cells met our thresholds for purity and enrichment scores to show significant co-clustering onto the mouse basal population, suggesting this population of cells in the adult fly is uniquely similar to mouse basal prostate cells. We next sought to examine the shared gene expression features between the fly main cells and mouse basal cells, hoping to reveal conserved, cell-type specific gene expression networks.

Complex biological datasets, such as scRNA-seq data, exhibit high dimensionality. This is due to the large number of features (e.g., thousands of genes, each with specific expression levels) measured across a relatively smaller number of samples (individual cells). This approach results in a vast amount of data: each gene represents a unique dimension, and with the capability to sequence thousands to millions of cells, researchers can accumulate extremely large datasets. However, a significant challenge in scRNA-seq data analysis is the sparsity of the datasets: many genes exhibit zero or near-zero expression in numerous cells, indicating they are not expressed.

This sparsity complicates the distinction between meaningful biological signals and noise, posing hurdles to comparative analysis. LIGER addresses these challenges through matrix factorization, specifically integrative non-negative matrix factorization (iNMF). It decomposes the gene expression matrix into three lower-dimensional matrices: the W-matrix (loading or feature weights matrix for shared signals), the V-matrix (loading or feature matrix for unshared signals within the data (e.g. batch effects, technical artifacts, etc.), and the H-matrix (scores or coefficients matrix). This decomposition, Data = H (W+V), effectively manages sparsity by identifying and reconstructing the non-zero values that represent actual gene expression, thereby revealing the data’s latent structure.

The W-matrix maps each gene to latent factors or "metagenes," inferred components that capture the dataset’s main sources of variation, such as cell types, states, or biological processes. This matrix is key for understanding how genes contribute to these metagenes, simplifying the data from tens of thousands of genes to a manageable number of factors. For example, a metagene strongly associated with cell cycle genes suggests variation in cell cycle states within the dataset. The H-matrix assigns each cell a score for these latent factors, providing insights into the cell-specific expression of the underlying biological processes. Together, these matrices enable researchers to navigate the complexity of scRNA-seq data, offering a clearer understanding of cellular heterogeneity and the mechanisms driving it.

LIGER allows for the mapping of W-matrix factors on UMAP clusters. We found that Factor 4 was enriched in the basal population (Figure 4A), suggesting the genes contributing to this metagene are actively expressed in the cells of that cluster and shared between fly main cells and mouse basal cells. Factor 4 contains 561 significant genes, consisting of genes that are co-expressed in both fly and mouse cells, as well genes that are expressed specifically by the mouse or fly cells that also contribute to main cell and basal cell co-clustering. The top 12 genes across the shared, fly-specific, and mouse specific lists are shown (Fig. 4 A, bottom).

**Figure 4:**
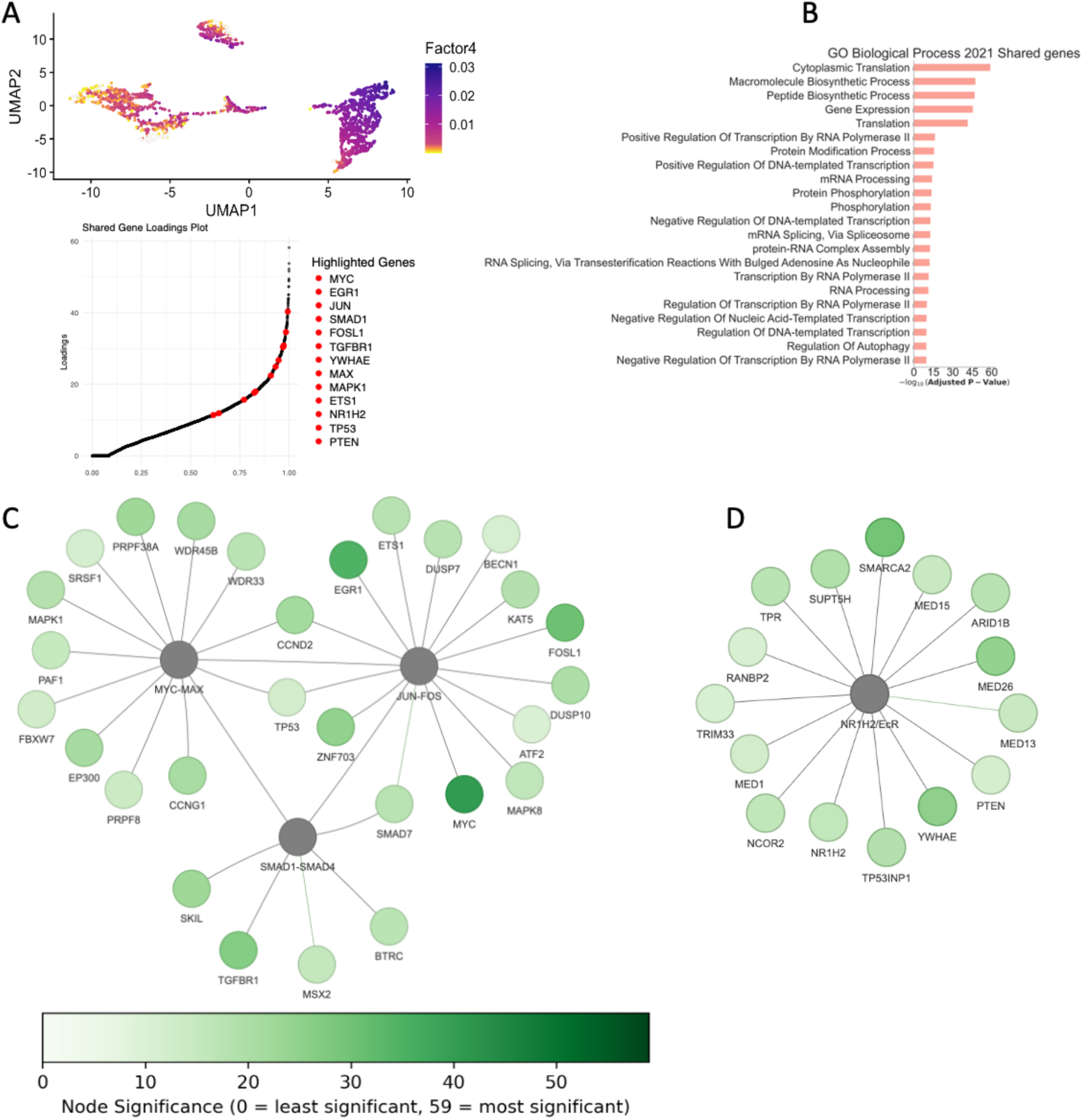
D*r*osophila Main Cells and Mouse Prostate Basal Cells have gene expression patterns consistent with Shared molecular Pathways A) Top: UMAP plot of the mouse/fly overlap indicating the loading strength of Factor4 metagene. Note Factor4 loading strength is high for the cluster where fly main cells and mouse basal cells co-cluster. Bottom: Top 12 genes in Factor4 metagene that are fly-specific, shared or mouse specific. Genes are sorted in descending order based on the magnitude of their contribution to factor loadings/metagene. B) GSEA plot of the top 22 GO-terms 561 for the LIGER Factor4 metagene shown in panel A. All genes had a fold change greater than 1 and an adjusted p-value less than 0.05. C) A Node graph representing the known connections between transcription factors in the shared gene set from Factor4 metagene and their known interactors or targets within the Factor4 metagene graphed with PyVis. D) A Node graph representing the known connections between the EcR transcription factors in the shared gene set from Factor4 metagene and targets genes from BioGrid, graphed with PyVis.

We focused on the shared set of genes in Factor 4 and performed Gene Set Enrichment Analysis (GSEA) for GO Biological Process (Subramanian et al. 2005). There was a large list of significant GO terms which we simplified and collapsed using ReviGO, which groups redundant GO terms (Supek et al. 2011). We find many GO terms enriched for processes expected to be engaged in highly secretory epithelial cells, such as mRNA transport, regulation of RNA processing, regulation of transcription initiation by RNA polymerase II, protein modification, synthesis, processing, and protein stability (Figure 4C). We also see the enrichment of genes involved in TOR signaling and stress-activated Ras/MAPK signal transduction. The RAS/MAPK pathway is found to be mis-regulated in primary and metastatic prostate lesions (Shen and Abate-Shen 2010; Wang et al. 2018; Taylor et al. 2010) and signaling via Ras/MAPK and

PI3K/AKT/TOR also induces tumorigenic-like phenotypes in the main cells of the *Drosophila* male accessory gland (Rambur et al. 2020).

Transcription factors that activate gene regulatory networks often reveal deep homology in cross-species comparisons (Shubin et al., 2009). We therefore pulled out the 44 conserved transcription factors in the shared Factor 4 gene set, which included several transcription factors known to play a role in prostate function, prostate growth and cell-type specific expression.

Notably this list includes JUN/FOSL1 (*kay/jra* in flies), MYC/MAX, and SMADs1 and 4 (*mad/medea* in flies), as well as the ETS transcription factor ETS1 (*pnt* or *Ets21C* in flies) (Fig 4C). This group of transcription factors is notable for their known roles in regulating gene expression and growth in the mammalian prostate as well as the *Drosophila* accessory gland (Church et al. 2025; Leiblich et al. 2019; Wang et al. 2018; Shen and Abate-Shen 2010).

However, they also regulate growth more broadly across many tissues. We therefore looked more closely at the other transcription factors that may be more cell-type specific such as the nuclear hormone receptors NR2C2, NR1H2, NR4A2 (*Hr78, EcR, Hr38* in flies respectively) and the androgen responsive FOXP1 (*Foxp* in flies) and the KLF family member KLF5 (*Dar1* in flies). Interestingly, KLF5 plays an essential role in mammalian prostate basal cells and is required in the basal bi-potential progenitors that give rise to both basal and luminal prostate cell types (Zhang et al. 2020). It will be interesting to examine roles for these transcription factors in the accessory gland to determine whether their targets in main cells and basal cells are conserved.

The Ecdysone receptor EcR has been shown to play a critical role in accessory gland growth and function and may serve a function analogous to Androgen Receptor (AR) signaling in the fly accessory gland (Leiblich et al. 2019; Sharma et al. 2017; Sekar et al. 2023). EcR target genes found in our analysis include regulators of transcription, such as subunits of the Mediator complex like MED1 and transcription cofactors like TRIM33 (fly gene name *bonus*) (Fig 4D).

MED1 is known to interact with several nuclear receptors, including AR, where its activity is increased in prostate cancer (PCa) and can mediate resistance to therapies (Russo et al. 2019). TRIM33 has been shown to promote the prostate cancer state by stabilizing AR, protecting it from Skp2-mediated ubiquitination and proteasomal degradation (Chen et al. 2022). The findings of shared expression profiles in genes known to be involved in prostate function and prostate cancer add additional credibility to the notion of a shared common ancestral cell type.

Our research has contributed to the understanding of similarities between the adult main epithelial cells of the *Drosophila* prostate-like accessory gland and the adult basal epithelial cells of the mammalian prostate. However, there remain additional questions to be addressed. Our analysis revealed overlaps in the transcriptional profiles of nuclear hormone receptor transcription factors. While the androgen receptor is the primary hormone receptor driving transcriptional changes in mammalian prostate cells, multiple hormone receptors may fulfill AR-like roles in *Drosophila*, with the steroid hormone ecdysone receptor being particularly significant. EcR is structurally similar to NR4A2. NR4A2 is involved in lipid metabolism as a sensor of cholesterol homeostasis in the mammalian prostate (Shiota et al. 2019). It is highly expressed in nonmalignant prostate epithelial RWPE-1 cells and normal human prostate epithelial cells but is significantly reduced in malignant-transformed prostate epithelial RWPE-2 cells and clinical prostate cancer samples (Shiota et al. 2019; Long et al. 2014). One role for EcR in *Drosophila* is to regulate lipid levels during pupation (Kamoshida et al. 2012), indicating a shared function with NR4A2, but EcR also has several other important roles during development. *Drosophila* AG models where RNAi to EcR or dominant negative forms of EcR are expressed in main cells lead to decreased fertility and dying cells, resulting in decreased levels of essential seminal proteins (Sharma et al., 2017). This highlights the importance of EcR function in the *Drosophila* prostate-like tissue.

Further research will be needed to determine the functional overlap between *Drosophila* nuclear receptors and mammalian nuclear receptors. Assuming a common prostate-like ancestral cell type, it is plausible that the patterns observed in *Drosophila* nuclear receptors evolved to specialize and diversify the signaling roles for mammalian nuclear receptors. Over evolutionary time, the many-faceted functions of EcR in hormone response and fertility in *Drosophila* may have been partially assumed by the mammalian androgen receptor (AR), while the lipid sensing-related functions were maintained by mammalian NR4A2.

There are additional questions regarding the other secretory epithelial cell type in the *Drosophila* accessory gland, the secondary cells. It is possible that both *Drosophila* main and secondary cells are transcriptionally similar to mammalian prostate basal cells. Previous research on mammalian prostate basal cells has shown that they express genes associated with stem cells, neurogenesis, and ribosomal RNA (rRNA) biogenesis (Zhang et al. 2016). While both the *Drosophila* main and secondary cells have secretory functions, in the case of main cells, the secretory function was not necessary for clustering with the mammalian prostate basal cell population based on transcriptional profiles. Alternatively, the transcriptional profile of *Drosophila* secondary cells may more closely parallel the mammalian prostate’s other epithelial cell type, the luminal cells. The Fly Atlas dataset used here did not contain enough secondary cells for our analysis pipeline, as there are 25 times more main cells than secondary cells per accessory gland lobe. As additional single-cell datasets of the *Drosophila* accessory gland become available, researchers can pool the data to achieve a sufficient number of secondary cells for analysis.

## Conclusion

Cross-species comparisons across large evolutionary distances present some unique challenges but also can reveal unappreciated similarities across cell types. Here we provide evidence of significant transcriptomic similarity between the adult main epithelial cells of the *Drosophila* prostate-like accessory gland and the adult basal epithelial cells of the mammalian prostate. These similarities could reflect shared evolutionary homology, or they could be independently derived features that emerge from shared tissue functions. Further work into the cell-type specific gene regulatory networks examining shared and unique features will be required to resolve these questions.

## Acknowledgements

We thank Dr. Mo Siddiq (Univ. Michigan) for valuable discussions regarding evolution and deep homology and Dr. Todd Morgan and Dr Aaron Udager (Univ. Michigan) for valuable discussions regarding prostate biology and prostate cellular markers. SJC was supported in part by NIH R35GM149273 to LB and a Rackham Merit Fellowship from the University of Michigan. ARK was supported by NIH F31HG012715. Work in the Welch Lab is supported by NIH R01HG010883. We used FlyBase (most recently release FB2025_02, released April 17, 2025) to find information on *Drosophila* gene sequences, phenotypes, and gene functions (Ozturk-Colak et al. 2024).

## Methods

**Software and Algorithms**
|  |  |  |
| --- | --- | --- |
| revigo | Supek et al., 2011 | http://revigo.irb.hr/index.jsp |
| LIGER | Welch et al., 2019 | https://github.com/welch-lab/liger |
| SCANPY | Wolf et al., 2018 | https://scanpy.readthedocs.io/en/stable/ |
| Matplotlib | (Hunter 2007) | https://matplotlib.org/ |
| Seaborn | (Waskom 2021) | https://seaborn.pydata.org/index.html |
| Pandas | McKinney et al., 2010 | https://pandas.pydata.org/ |
| Gseapy | Fang et at., 2023 | https://gseapy.readthedocs.io/ |
| DIOPT | Hu et al., 2011 | https://fgr.hms.harvard.edu/diopt |
| Pyvis |  | https://pyvis.readthedocs.io/en/latest/ |

### DIOPT

We obtained a list of fly genes from FlyBase, using "bgn_annotation_ID" and "<u>gene_snapshots_fb_2023_06</u>". This gene list was input into the <u>DIOPT version 9</u> database to find homologous gene pairs. We applied a similar approach to genes identified in the mouse prostate dataset. Homologous gene pairs were filtered as described previously. Genes with orthologs in both fly and mouse datasets were assigned human names. Otherwise, fly genes were appended with ‘_D’ and mouse genes with ‘_M’ to ensure none identified genes were not incorporated into the similar features in both datasets.

### Figures and graphs

Figure 1A was created with Dall-E 3 and PowerPoint. Figure 1B-G, Figure 2G, and Figure 3C were generated via Seaborn and Matplotlib. Figure 2A-C were generated via SCANPY. Figure 2D-F, Figure 3A-B, and Figure 4A were generated via LIGER. Figure 4B was generated via GSEApy. Figure 4C was generated by manually providing target and edge nodes from our significant genes and visualizing with the pyvis.network.

### Liger and SCANPY Workflow

We followed the workflow described in <u>LIGER</u> and <u>SCANPY</u> webpages for

1. Loading dataset to create LIGER or AnnData object
2. Preprocessing, normalization, and removing low quality cells.
3. In SCANPY we used minimum gene threshold of 500 and transcripts expressed in a minimum of 3 cells.
4. Joint Matrix Factorization for LIGER or Principal component analysis for SCANPY
5. Quantile Normalization, Computing the neighborhood graph, and Joint Clustering
6. With LIGER we use a variable threshold setting between 0.1 - 0.5 depending on the fly tissue being compared to the mouse basal populations. Optimizing for mouse cell purity. We started with a λ = 5, K = 30 for all tissues comparison. Then moved on to using the built-in optimizeNewK and optimizeNewLambda features of LIGER for values ranging from λ = 5-10 and K = 5-30, optimizing for mouse cell purity and fly cell overlap with mouse basal population.
7. LIGER used Louvain clustering, and SCANPY used Leiden.
8. Mouse cells used a n_neighbors setting of 10 and n_pcs of 40 for clustering the mouse populations.

2. Visualization via UMAPS

---

**Supplemental Figure 1.**
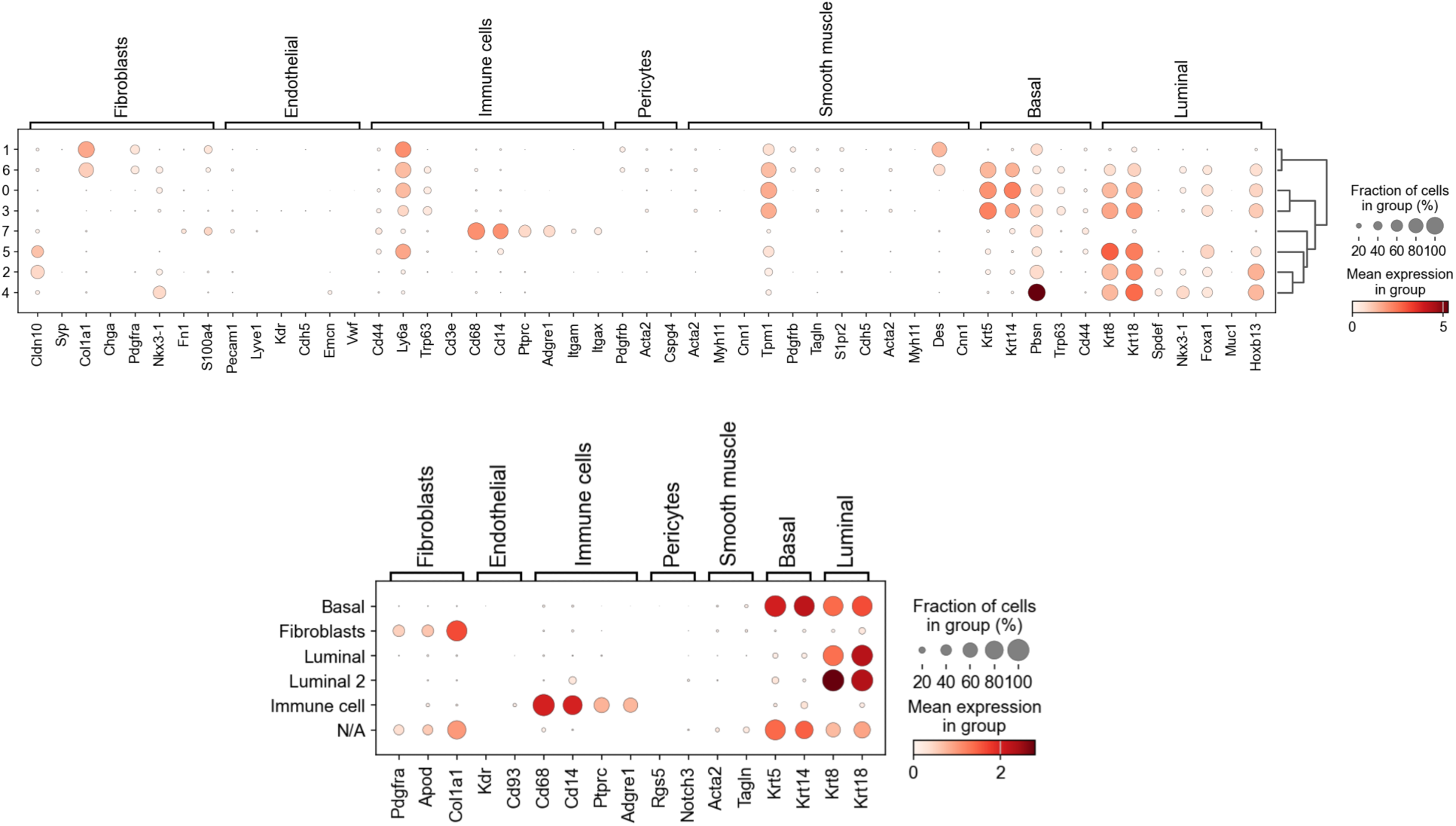
Markers for prostate cell types. (Top) The full gene list curated from the Human Protein Atlas and for assignment of mouse prostate cell types to clusters 0-7. (Bottom) the smaller marker list used to annotate broad mouse prostate cell types in Fig. 2A.

